# Temporal cascade of frontal, motor and muscle processes underlying human action-stopping

**DOI:** 10.1101/700088

**Authors:** Sumitash Jana, Ricci Hannah, Vignesh Muralidharan, Adam R. Aron

**Affiliations:** Department of Psychology, University of California, San Diego

## Abstract

Action-stopping is a canonical executive function thought to involve top-down control over the motor system. Here we aimed to validate this stopping system using high temporal resolution methods in humans. We show that, following the requirement to stop, there was an increase of right frontal beta (∼13 to 30 Hz) at ∼120 ms, likely a proxy of right inferior frontal gyrus; then, at 140 ms, there was a broad skeletomotor suppression, likely reflecting the impact of the subthalamic nucleus on basal ganglia output; then, at ∼160 ms, suppression was detected in the muscle, and, finally, the behavioral time of stopping was ∼220 ms. This temporal cascade confirms a detailed model of action-stopping, and partitions it into subprocesses that are isolable to different nodes and are more precise than the behavioral speed of stopping. Variation in these subprocesses, including at the single-trial level, could better explain individual differences in impulse control.

---

The ability to control one’s actions and thoughts is important for our daily lives; for example: changing gait when there is an obstacle in the path^1^, resisting the temptation to eat when on a diet^2^, overcoming the tendency to say something hurtful^3^. While many processes contribute to such forms of control, one important process is response inhibition – the prefrontal (top-down) stopping of initiated response tendencies^4^. In the laboratory, response inhibition is often studied with the stop-signal task^5^. On each trial, the participant initiates a motor response, and then, when a subsequent Stop signal occurs, tries to stop. From the behavioral data one can estimate a latent variable; the speed of stopping known as Stop Signal Reaction Time (SSRT), which is typically 200-250 ms in healthy adults^5^. SSRT has been useful in neuropsychiatry where it is often longer for patients vs. controls^6–11^. The task has also provided a rich test-bed, across species, for mapping out a putative neural architecture of prefrontal-basal-ganglia-regions for rapidly suppressing motor output areas^6,12,13^. Given this rich literature, this task is one of the few paradigms included in the longitudinal Adolescent Brain Cognitive Development study^14^ of 10,000 adolescents over 10 years.

Against this background, a puzzle is that the relation between SSRT and ‘real-world’ self-reported impulsivity is often weak^15–20^. One explanation is that SSRT may not accurately index the brain’s true stopping speed. Indeed, recent mathematical modelling of behavior during the stop-signal task suggests that standard calculations of SSRT may overestimate the brain’s stopping speed by ∼100 ms^15^ [also see^21^]. Further, in a recent study^22^, electromyographic (EMG) recordings revealed an initial increase in EMG activity in response to the Go cue, followed by a sudden decline at ∼150 ms after the Stop signal. This decline in EMG could be because of the Stop process ‘kicking in’ to cancel motor output – but the striking thing is that this was 50 ms before the SSRT of 200 ms. This timing is also consistent with experiments using transcranial magnetic stimulation (TMS) to measure the motor evoked potential (MEP) during the stop-signal task (the MEP indexes the excitability of the pathways from motor cortex to muscle). The MEP in the muscle that was-to-be-stopped reduced at ∼150 ms^23,24^. Further, other studies that measured the MEP from muscles that were not needed for the task, show there is ‘global suppression’ also at ∼150 ms^25–28^ (*i.e.* corticospinal activity was suppressed for the broader skeletomotor system). This ‘global MEP suppression’ has been linked to activation of the subthalamic nucleus of the basal-ganglia^29^, which is thought to be critical for stopping, and might broadly inhibit thalamocortical drive^30^.

The potential overestimation of the brain’s true stopping speed by SSRT could arise for several reasons. First, the race model assumes that the Stop process is “triggered” on every trial. But recent research shows that this is not the case^15^, and that failing to account for “trigger failures” inflates SSRT. Second, while the standard “race model” assumes that the Go and Stop processes are independent^5^, recent research show that violations of this independence can also inflate SSRT^21^. Finally, the standard ways of computing SSRT likely do not account for electromechanical delays between muscle activity and the response. In any event, overestimating the brain’s stopping speed would add variance to SSRT which could potentially weaken the above-mentioned across-participant associations between stopping speed and self-report scores^15–17^. Furthermore, if the true stopping speed is ∼150 ms, the timing of activation of nodes in the putative response inhibition network should precede this time-point for those nodes to play a causal role in action stopping – and this is important for the interpretation of neuroscience studies. For instance, in electrocorticography and electroencephalography (EEG) studies, successful stopping elicits increased beta band power over right frontal cortex in the time period between the Stop signal and SSRT^31–33^. Whether this, and other, neurophysiological markers of the Stop process occur sufficiently early to directly contribute to action-stopping (if SSRT is overestimated) is unknown; yet this is fundamental to our understanding of brain networks underlying response inhibition.

Here we leveraged the insight from the above-mentioned study^22^ which used EMG of the task relevant muscles. We now tested whether we could derive a single trial estimate of stopping speed from EMG (referred to as CancelTime). More specifically, we hypothesized that ‘partial’ EMG bursts on the Successful Stop trials (*i.e.* small EMG responses that begin but do not reach a sufficient amplitude to lead to an overt response)^34^ would carry information about the latency of stopping and tested this in two studies. In a third study we tested if CancelTime would correspond with the measure of putative basal ganglia-mediated global motor suppression, measured with single-pulse TMS. In studies four and five we turned to the cortical process thought to initiate action–stopping, using the above-mentioned proxy of right frontal beta^31,32^. We measured scalp EEG, derived a right frontal spatial filter in each participant, and then extracted beta bursts^35^ in the time period between the Stop signal and SSRT. We tested how the timing of these beta bursts related to CancelTime.

## Results

### Study 1 (EMG)

10 participants performed the stop-signal task (Fig. 1a). On each trial they initiated a manual response when a Go cue occurred, and then had to try to stop when a Stop signal suddenly appeared on a minority of trials. Depending on the stop signal delay, SSD, participants succeeded or failed to stop, each ∼50% of the time). We measured EMG from the responding right index and little fingers (Fig. 1b *inset*). Behavioral performance was typical, with SSRT (referred to as SSRT_Beh_) of 216±8 ms, and action-stopping on 51±1 % of Stop trials (Table 1). EMG analysis was performed on the trial-by-trial root-mean-squared EMG (EMG_RMS_; Fig. 1b). On 53±6 % of Successful Stop trials (*i.e.* where no keypress was made) there was a small but detectible EMG response (Partial EMG trials; see Supplementary Fig. 1a, b for EMG-RT correlation), while on the remainder of Successful Stop trials there was no detectible EMG response (No EMG trials). The amplitude of EMG responses (mean peak EMG voltage) in the Partial EMG trials was 48±3% smaller than in trials with a keypress (Fig. 2a).

**Figure 1.**
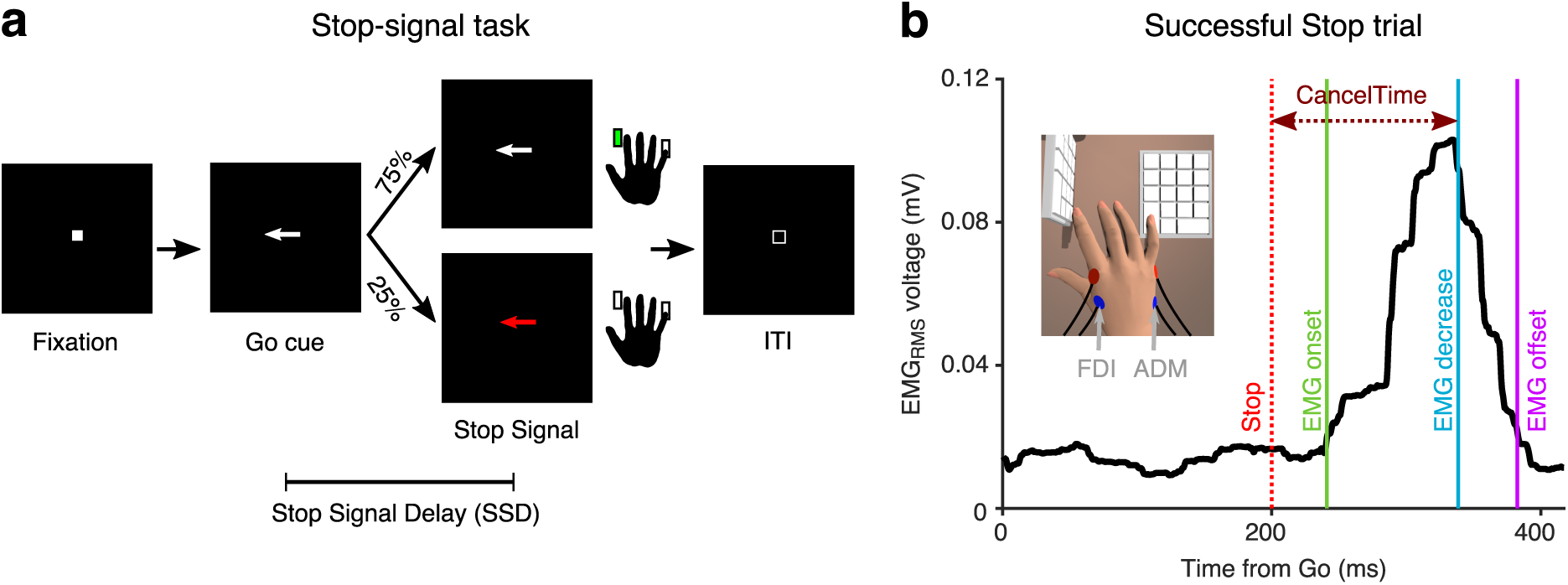
Behavioral task and EMG recording. (**a**) Stop-signal task. (**b**) EMG_RMS_ on a Successful Stop trial (Partial EMG) in an exemplar participant, aligned to the Go cue. CancelTime refers to the time from the Stop signal (dotted red line) to when the EMG_RMS_ starts decreasing (blue line). The green and purple line represent the detected onset and offset of the EMG response. (*Inset*) Recording set-up with a vertical and a horizontal keypad to record keypresses from the FDI and ADM muscles.

**Figure 2.**
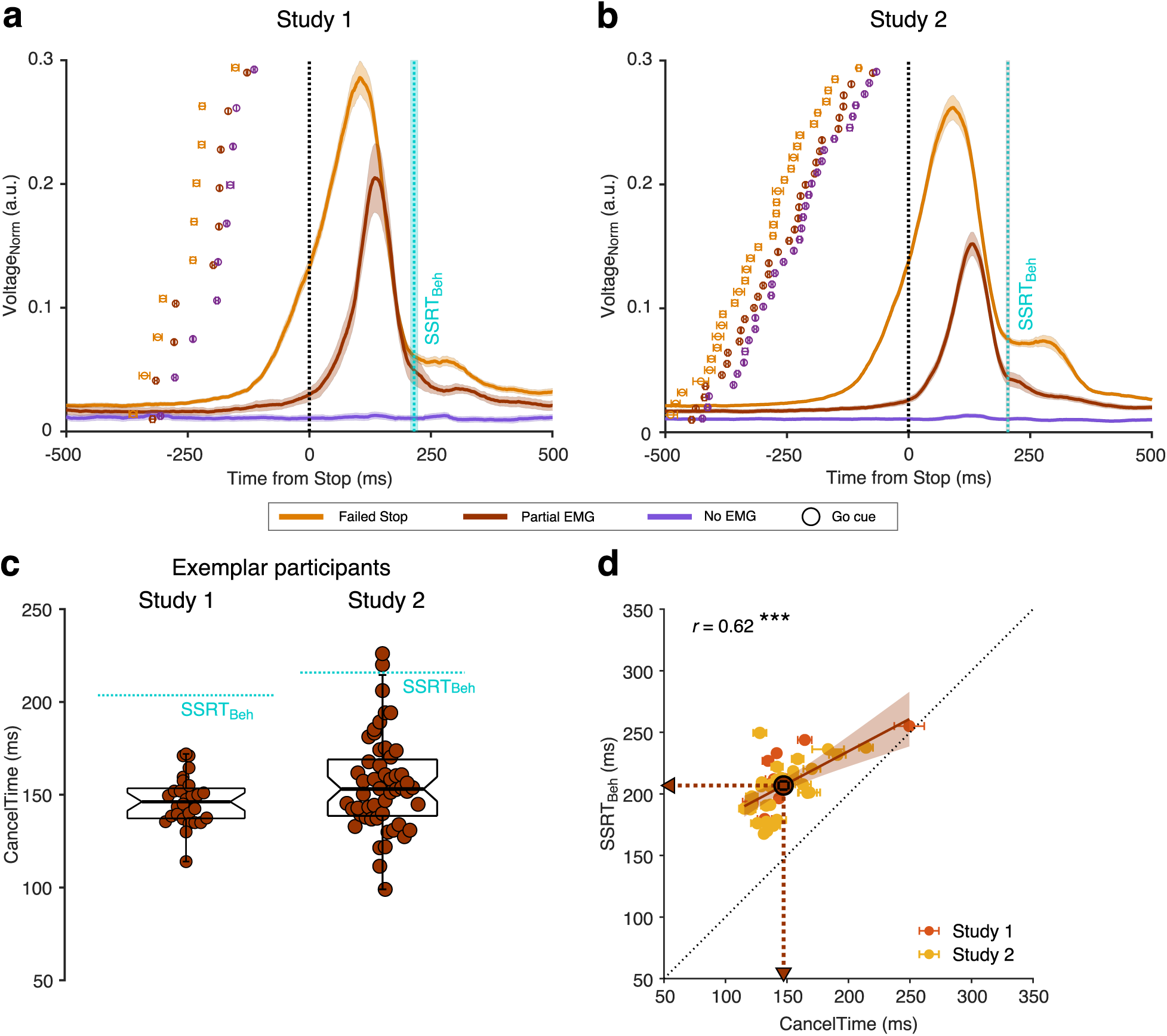
EMG responses in study 1 and 2. (**a**) Normalized EMG_RMS_ voltage in Failed Stop (orange), Partial EMG (brown), and No EMG trials (purple), aligned to the Stop signal. The lines and the shaded area represent the mean±s.e.m. across participants. The dotted cyan line and shaded area represent the mean±s.e.m of SSRT_Beh_ across participants. The dots and cross-hairs represent the mean±s.e.m. of the Go cue in a participant. Note that the time between the Go cue and the Stop signal (i.e. the SSD) is shortest for the No EMG (purple), then the Partial EMG (brown), and then the Failed Stop trials (orange). (**b**) Same as (a) but for Study 2. (**c**) (*Right*) Beeswarm plot of the CancelTime in an exemplar participant from Study 1. Each dot represents a trial. The dotted cyan line represents the SSRT_Beh_. (*Left*) Same as *right* but for Study 2. (**d**) Correlation between CancelTime and SSRT_Beh_ in Study 1 (light red) and Study 2 (yellow). The brown dot, lines and arrows represent the means, while the black dotted line represents the unity line. The linear regression fit and its 95% confidence interval (pooled study 1 and 2) is shown as a brown line and shaded region respectively.

**Table 1:** Behavior. (mean±s.e.m.; All values in ms)

|  | Study 1<br>(EMG) | Study 2<br>(EMG) | Study 3<br>(TMS) | Study 4<br>(EEG) | Study 5<br>(EEG) |
| --- | --- | --- | --- | --- | --- |
| **Go RT** | 470 (15) | 493 (15) | 430 (17) | 427 (15) | 405 (6) |
| **Failed Stop RT** | 416 (11) | 447 (14) | 391 (12) | 384 (12) | 370 (5) |
| **Correct Go %** | 97 (1) | 98 (0) | 99 (0) | 99 (0) | 99 (0) |
| **Correct Stop %** | 51 (1) | 52 (1) | 49 (1) | 48 (1) | 50 (0) |
| **Mean SSD** | 237 (20) | 280 (17) | 194 (18) | 191 (21) | 170 (7) |
| **SSRT<sub>Beh</sub>** | 216 (8) | 204 (4) | 219 (6) | 214 (9) | 219 (6) |

We hypothesized that the time when the Partial EMG response starts declining after the Stop signal is a readout of the time when the Stop process is implemented in the muscle (hereafter ‘CancelTime’). We observed that, first, CancelTime is much earlier than SSRT_Beh_ (see Fig. 2c (*left*) for all CancelTimes in an exemplar participant; mean CancelTime = 146±3 ms, SSRT_Beh_ = 203 ms); and second, across participants, CancelTime was positively correlated with SSRT_Beh_ (Fig. 2d; study 1: mean CancelTime = 152±11 ms, mean SSRT_Beh_ = 216±8 ms; *r* = 0.71, *p* = 0.020, *BF_10_* = 3.6). This suggests that CancelTime might index the time when Stopping is implemented at the muscle.

### Study 2 (EMG)

We then ran a new sample (*n* = 32; see Table 1 for behavioral results). Again, we observed partial EMG responses on 49±2 % of Successful Stop trials; where the EMG amplitude was 54±1 % smaller than the amplitude in trials with a keypress (Fig. 2b). Fig. 2c (*right*) shows the distribution of CancelTimes in an exemplar participant (mean CancelTime = 156±4 ms, SSRT_Beh_ = 218 ms). Again, across participants, mean CancelTime was positively correlated with SSRT_Beh_ (Fig. 2d; mean CancelTime = 146±4 ms, mean SSRT_Beh_ = 204±4 ms; *r* = 0.59, *p* < 0.001, *BF_10_* = 71.7). Intriguingly, in each study, CancelTime was ∼60 ms less than SSRT_Beh_. To further explore this, we pooled the data across the 2 studies.

### Pooled studies 1 and 2

Mean CancelTime (147±5 ms) was 60±3 ms shorter than SSRT_Beh_ (*t*(41) = 18.4, *p* < 0.001, *d* = 2.5, *BF_10_* > 100). We then tested whether we could calculate SSRT using the presence of EMG responses (SSRT_EMG_) instead of the keypress responses (SSRT_Beh_). We considered Partial EMG trials as Failed Stop trials and used EMG onset time on Correct Go trials to recalculate SSRT (*i.e.* instead of using P(Respond|Stop) from behavior and Go RT as is typical for SSRT_Beh_ calculations; see Methods; see Fig. 3a for an exemplar participant). We then performed 1-way repeated measures ANOVA with “Stop Time” as the dependent measure and the method of estimation as a factor (SSRT_EMG_, SSRT_Beh_, and CancelTime). There was a significant main effect of the estimation method on “Stop Time” (*FGG*(1.4, 56.1) = 66.3, *p* < 0.001, 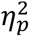 = 0.6). Pairwise comparisons showed that SSRT_EMG_ (157±7 ms) was significantly faster than SSRT_Beh_ (209±3 ms) (Fig. 3b; *t*(41) = 8.2, *p_Bon_* < 0.001, *d* = 1.3, *BF_10_* > 100), but importantly, not significantly different from mean CancelTime (*t*(41) = 1.5, *p_Bon_* = 0.270, *d* = 0.2, *BF_10_* = 0.5). This suggests that SSRT_Beh_ might be protracted by a peripheral delay and that CancelTime might be a better metric of the time of implementation of the Stop process.

**Figure 3.**
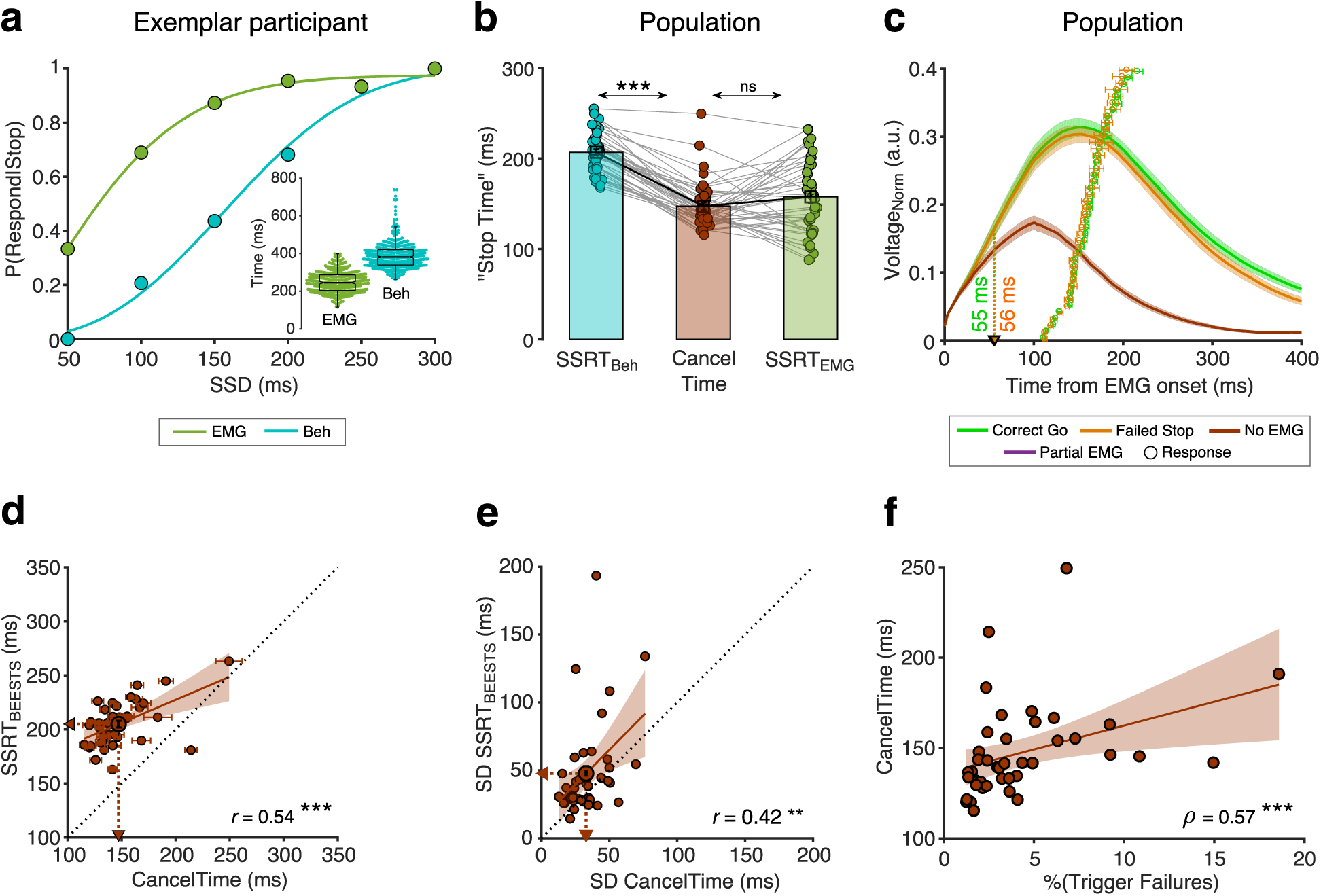
Peripheral delay associated with SSRT_Beh_ and relationship between CancelTime and BEESTS parameters. (**a**) P(Respond|Stop) in an exemplar participant calculated using the behavioral response (dark green dots) and the EMG response (cyan dots). The lines represent the cumulative Weibull fit as *W*(*t*) = *γ* – (*γ* – *δ*)*e*^[−(*t/α*)*^β^*]^ where where *t* is the SSD, *α* is the time at which the function reaches 64% of its full growth, *β* is the slope, *δ* is the minimum value of the function, and *γ* is maximum value of the function. The difference between *δ* and *γ* marks range of the function. (*Inset*) Beeswarm plot of the EMG onset (dark green) and the behavioral responses (cyan) used to calculate SSRT_EMG_ and SSRT_Beh_ respectively. (**b**) Comparison of the SSRT_Beh_ (cyan), CancelTime (brown), and SSRT_EMG_ (dark green) across all participants. Each dot represents a participant, while the bar and cross-hair represents the mean±s.e.m. in a group. (**c**) The normalized EMG responses aligned to the detected EMG onsets in the Correct Go (dark green), Failed Stop (orange), and Partial EMG (brown) trials. The line and shaded region represent the mean±s.e.m. in a group. The dots and cross hairs represent the mean±s.e.m. of the keypress in a participant. (**d**) Correlation between CancelTime and mean SSRT_BEESTS_ estimate. Each dot and cross-hair represent the mean±s.e.m. in a participant. The brown line and the shaded area represent the linear regression fit and its 95% confidence interval. The unity line is represented as a dotted black line. (**e**) Correlation between CancelTime and SD of the SSRT_BEESTS_ estimate. Other details same as (d). (**f**) Correlation between percentage Trigger Failures estimated from BEESTS and CancelTime. Other details same as (d).

Next, we examined in more detail the EMG profile on Partial EMG trials. Across all participants, the EMG response in the Partial EMG trials (when aligned to the EMG onset) had a profile similar to the EMG response in the Correct Go and Failed Stop trials, but diverged ∼55 ms after EMG onset (55 ms compared to Correct Go, and 56 ms compared to Failed Stop trials, Fig. 3c). We surmised that if the Partial EMG trials reflect responses that have been actively cancelled at the muscle-level, then the amplitude of these responses should increase with SSD. The rationale was that, at shorter SSDs, the Go process will have been active for a shorter duration, meaning EMG activity will not have increased much before being inhibited, while at longer SSD, the Go process will have been active for a longer duration, meaning EMG activity will have increased much more before being inhibited. Indeed, the amplitude of the Partial EMG responses increased with SSD (Supplementary Fig. 1c). A 1-way repeated measures ANOVA with amplitude as the dependent variable and the SSD as the independent variable showed significant effect of SSD on amplitude (*F*(4,24) = 3.7, *p* = 0.018, 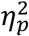 = 0.4) [also see^23^]. This suggests that the Partial EMG trials represent inhibited Go responses and not merely a weak Go process (which would presumably not increase across SSDs).

To further validate CancelTime, we modelled the behavior using BEESTS (Bayesian Estimation of Ex-gaussian STop-Signal reaction time distributions; see Table 2 for model estimates). While SSRT_Beh_ produces a single estimate per person, BEESTS uses a Bayesian parametric approach to estimate the distribution of SSRTs^36^. Also, for each participant, it provides an estimate of the probability of trigger failures (*i.e.* stop trials where the stopping process was not initiated^36^). Across participants, mean CancelTime was positively correlated with the mean SSRT_BEESTS_ (205±3 ms; *r* = 0.54, *p* < 0.001, *BF_10_* > 100; Fig. 3d). More interestingly, the SD of CancelTime (33±2 ms) was positively correlated with the SD of SSRT_BEESTS_ (48±5 ms; *r* = 0.42, *p* = 0.005, *BF_10_* = 6.9; Fig. 3e). Further, the percentage of trigger failures (4±1%) was positive correlated with mean CancelTime (*ρ* = 0.57, *p* < 0.001, *BF_10_*> 100) suggesting that participants who fail to “trigger” the Stop process more often, have longer CancelTimes (Fig. 3f). These relationships between CancelTime and model estimates give further credence to our interpretation that CancelTime on Partial EMG trials reflects a single-trial measure of the time of implementation of the Stop process.

**Table 2:**
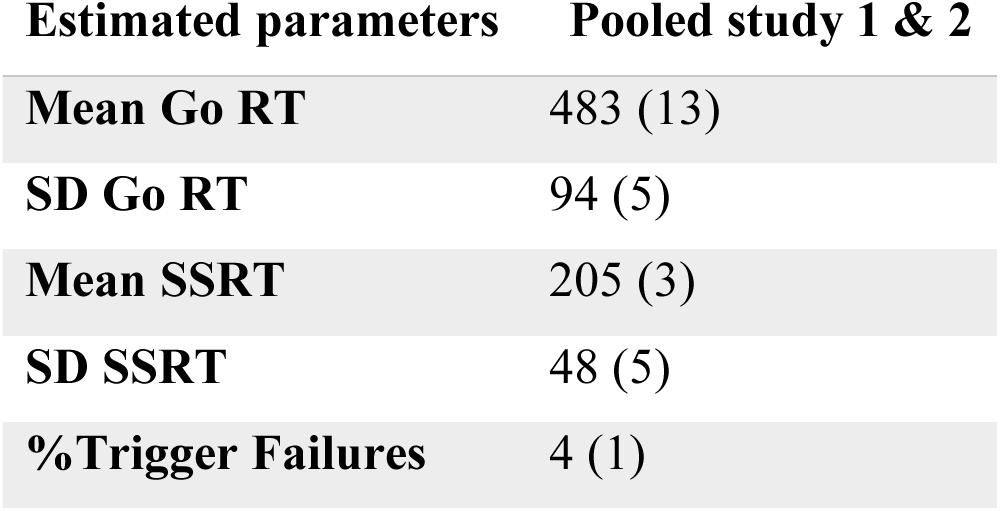
BEESTS estimates. (mean±s.e.m.; All values in ms)

| Estimated parameters | Pooled study 1 & 2 |
| --- | --- |
| **Mean Go RT** | 483 (13) |
| **SD Go RT** | 94 (5) |
| **Mean SSRT** | 205 (3) |
| **SD SSRT** | 48 (5) |
| **%Trigger Failures** | 4 (1) |

### Study 3 (TMS)

To further validate CancelTime and relate it to brain processes we turned to a different method – single-pulse TMS over a task-irrelevant muscle representation in the brain. As mentioned above, the reduction of MEPs from task-irrelevant muscles on Successful Stop trials^25–27^, is thought to reflect a basal ganglia-mediated global suppression^29^. Eighteen new participants (see Table 1 for behavioral results) now performed the task with their left hand, while TMS was delivered over the left motor cortex and MEPs were recorded from a task-irrelevant, right forearm muscle. MEPs were recorded at different times after the Stop signal on different trials: 100 – 180 ms in 20 ms intervals, as well as during the inter-trial interval which served as a baseline. Concurrently, we recorded EMG from the task-relevant left-hand muscles as for studies 1 and 2 above (Fig. 4a).

**Figure 4.**
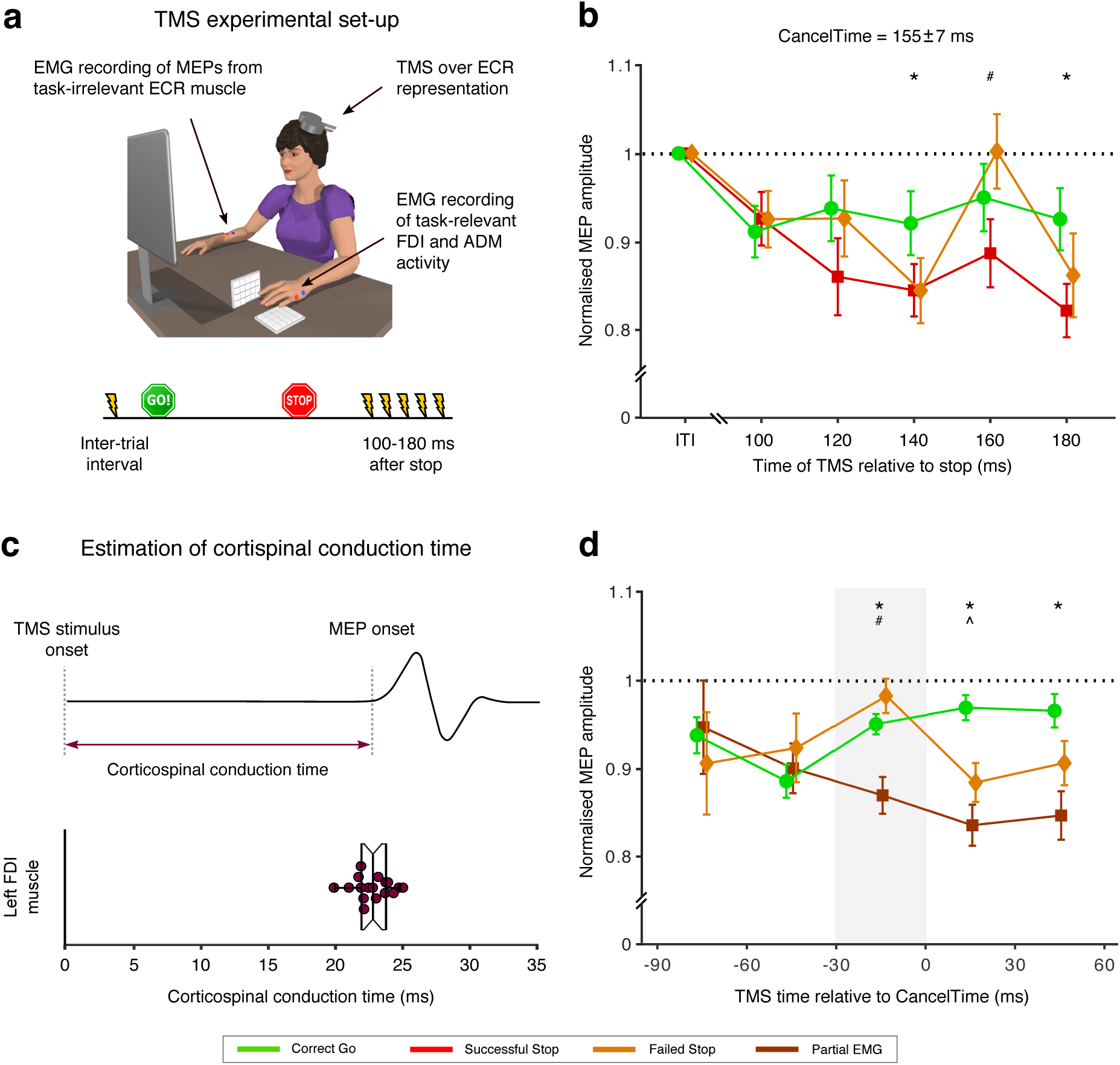
Relationship between global motor system suppression and CancelTime. (**a**) Experimental set up and TMS stimulus timings for study 3. Participants performed the Stop signal task with the left hand with concurrent EMG measurement of CancelTime from task-relevant muscles FDI and ADM muscle. On a given trial, a single TMS stimulus over left M1 was delivered at one of 6 possible times to elicit a motor evoked potential (MEP) in the task-irrelevant extensor carpi radialis (ECR) muscle of the right forearm. (**b**) Global motor system suppression begins at 140 ms after the Stop signal, and thus ∼15 ms prior to the mean CancelTime. Paired *t-*tests: *\*, p_Bon_* < 0.05 Successful Stop (red) vs. Correct Go (green); *#*, *p_Bon_* < 0.05 Successful Stop vs. Failed Stop (orange). The black dotted line shows amplitude of MEPs normalized to those at the inter-trial interval. (**c**) (*Top*) Schematic representation of an MEP. (*Bottom*) Beeswarm plot of the mean corticospinal conduction time to a hand muscle was established by measuring the onset latency of MEPs in the hand, and was ∼23 ms on average. Each dot represents a participant. This conduction time is included in CancelTime. (**d**) Trial-by-trial analysis of MEP amplitudes organized into 30 ms time bins reflecting the time of TMS expressed relative to the CancelTime. Global motor system suppression begins in a window 30-0 ms prior to the CancelTime (gray shaded region). Wilcoxon rank sum test: *\*, p_Bon_* < 0.05 Partial EMG (brown) vs. Correct Go (green); *#*, *p_Bon_* < 0.05 Partial EMG vs. Failed Stop (orange); ^, *p_Bon_* < 0.05 Failed Stop vs. Correct Go. The black dotted line shows amplitude of MEPs normalized to those at the inter-trial interval.

The key TMS finding, in keeping with earlier studies^25–27^, was of suppression of MEPs in the task-irrelevant forearm, indicating global motor system suppression, beginning ∼140 ms following the Stop signal in Successful Stop trials (Fig. 4b) [see Supplementary Fig. 2 for MEP amplitudes for Partial EMG and No EMG trials separately]. A 2-way repeated measures ANOVA with MEP amplitude as the dependent measure and the factors of trial-type (Correct Go, Successful Stop, Failed Stop) and time (100, 120, 140, 160, 180 ms after the Stop signal) showed main effects of both trial-type (*F*(2,32) = 7.2, *p* = 0.003, *η_p_^2^*= 0.3) and time (*FGG*(2.5, 40.7) = 4.8, *p* = 0.008, *η_p_^2^* = 0.2), as well as an interaction of trial-type by time (*F*(8, 128) = 3.4, *p* = 0.002, *η_p_^2^* = 0.2). Post hoc *t*-tests across Successful Stop and Correct Go trials showed *no* difference at 100 ms (*t*(16) = 0.7, *p_Bon_* = 1.0, *BF_10_* = 0.3), 120 ms (*t*(16) = 2.5, *p_Bon_*= 0.066, *BF_10_* = 2.8), and 160 ms (*t*(16) = 2.1, *p_Bon_*= 0.159, *BF_10_* = 1.4). However, MEP amplitudes *were* significantly suppressed on Successful Stop trials at 140 ms (*t*(16) = 4.1, *p_Bon_*= 0.003, *BF_10_* = 39.8) and 180 ms (*t*(16) = 4.4, *p_Bon_*< 0.001, *BF_10_* = 65.2) after the Stop signal. Therefore, we estimate the onset of the global motor suppression to be ∼140 ms after the Stop signal, which places it ∼15 ms prior to the mean CancelTime (155±7 ms). There were no significant differences in MEP amplitudes between Failed Stop and Correct Go trials at any time point, though MEP amplitudes on Successful Stop trials were also suppressed compared to Failed Stop trials at 160 ms (*t*(16) = 2.9, *p_Bon_* = 0.033, *BF_10_* = 4.9).

It makes sense that global motor suppression occurs before CancelTime as motor cortical output takes time to be transmitted along the corticospinal pathway to the muscles. To verify whether the ∼15 ms discrepancy in timings could be accounted for by corticospinal conduction delays, we estimated this corticospinal conduction time in a separate phase of the current study by delivering TMS over the hand representation to evoke MEPs in the left, task-relevant, FDI muscle (Fig. 4c). This was 23± 0.3 ms. Thus, a decline in muscle activity would be expected to be preceded by a reduction in motor cortical output by ∼23 ms, which is very similar to the ∼15 ms difference between global motor suppression and CancelTime.

To further elaborate the temporal relationship between global motor suppression and CancelTime, we performed a trial-by-trial analysis whereby MEP amplitudes were sorted according to the time at which TMS was delivered, relative to the time at which EMG decreased on Successful Stop, Failed Stop and Correct Go trials (Fig. 4d). The suppression of MEPs in Successful Stop trials compared to Correct Go trials began in the 30 ms prior to the EMG decline (−30 to 0 ms: *Z* = 3.12, *p_Bon_* = 0.005; 0 to 30 ms: *Z* = 4.48, *p_Bon_* < 0.001; 30 to 60 ms: *Z* = 2.45, *p_Bon_* = 0.045). This lag in the time of EMG decrease relative to the time of the MEP suppression on Successful Stop trials can again be accounted for by the corticospinal conduction time. Thus, these results imply that the brain output to task-relevant muscles declines at approximately the same time as the global motor suppression begins.

### Study 4 (EEG)

Having established that CancelTime reflects the time of an active stopping process at the muscle (Studies 1 and 2, EMG/behavior), which also related tightly with the timing of global motor suppression (Study 3, TMS), we then tested whether this EMG measure was also related to the timing of a prefrontal correlate of action-stopping, specifically the increase of beta power (13-30 Hz) before SSRT_Beh_ at right frontal electrode sites^32,33^. We now measured scalp EEG as well as EMG from the hand, in 11 participants (see Table 1 for behavioral results). We derived beta bursts rather than beta power *per se*, as bursts have richer features^37^, such as burst timing and duration.

To identify right frontal electrodes of interest in each participant (*i.e.* a spatial filter), we used Independent Components Analysis^38^ [see^32,33^]. We selected a participant-specific independent component (IC) based on two criteria; First, the scalp topography (right-frontal, and if not present, frontal); and Second, an increase in beta power on Successful Stop trials (from Stop signal to SSRT_Beh_; Stop_Win_) compared to activity *prior to the Go* cue [-1000 to −500 ms aligned to the Stop signal; see Methods; Supplementary Fig. 3]. The average scalp topography across all participants is shown in Fig. 5b *inset*. For each participant, we estimated beta bursts; First, by filtering the data at the peak beta frequency; and Second, by defining a burst threshold based on the beta amplitude in a baseline period *after* the Stop signal (500-1000 ms after Stop signal in the Stop trials, and 500-1000 ms after the mean SSD in the Correct Go trials) (see Methods; Supplementary Fig 4).

**Figure 5.**
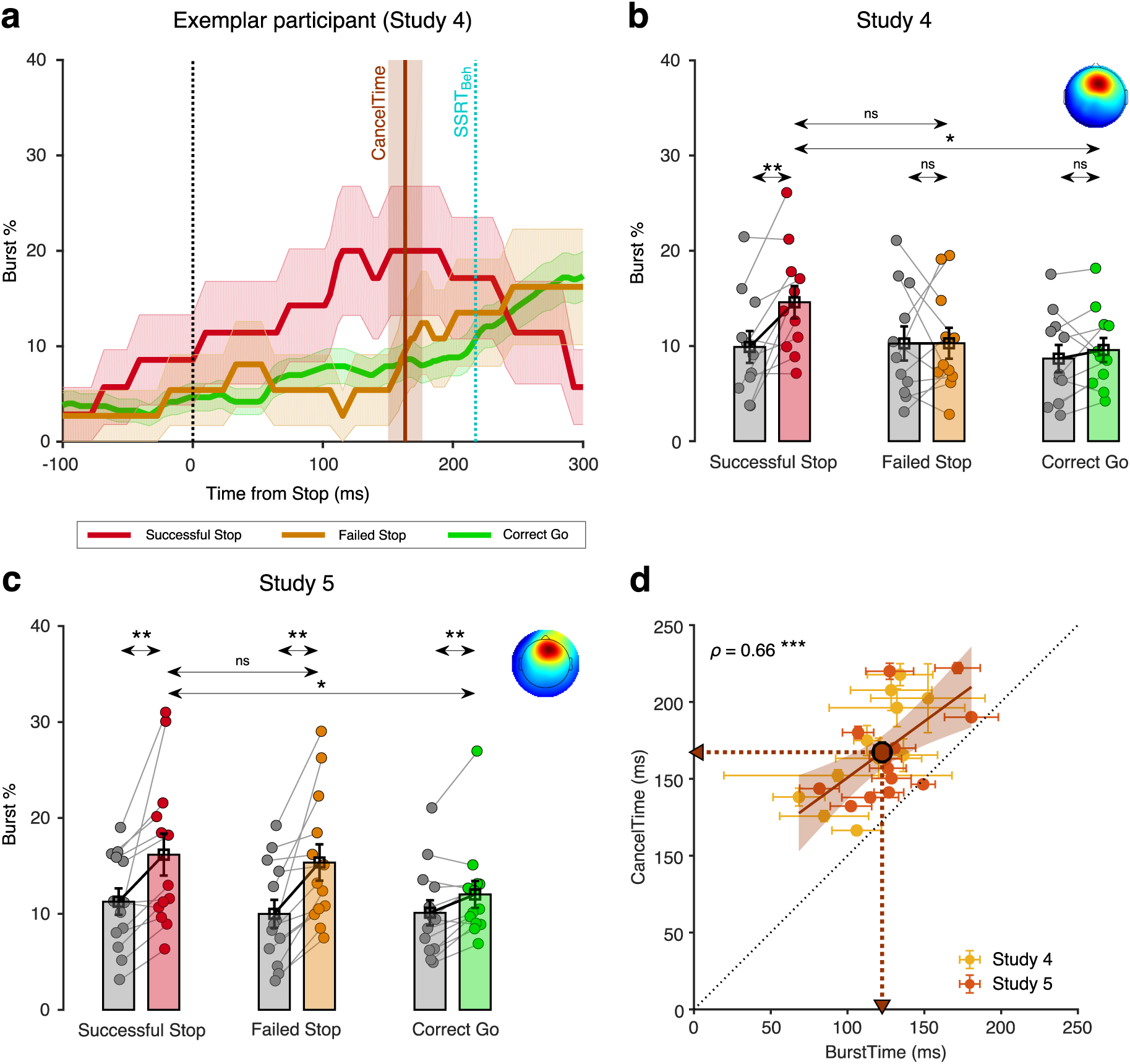
Relationship between scalp EEG beta bursts and CancelTime (study 4 and 5). (**a**) Burst % across time for Successful Stop (red), Failed Stop (orange), and Correct Go (green) trials for an exemplar participant in study 4 from the right frontal spatial filter. The shaded region represents mean±s.e.m. The CancelTime is shown in brown and the SSRT_Beh_ as a cyan line. (**b**) The mean burst probability across all participants for Successful Stop (red), Failed Stop (orange), and Correct Go (green) trials and their respective baselines (gray). The bars and cross-hairs represent the mean and s.e.m across participants, while the dots represent individual participants. (*Inset top right*) The average scalp topography of all the right frontal ICs across all participants. (**c**) Same as (b) but for study 5. (**d**) Correlation between mean BurstTime and mean CancelTime. The yellow dots and cross-hairs represent the participants in study 4, while the light red ones represent participants in study 5. The brown line and the shaded area represent the linear regression fit and its 95% confidence interval (pooled study 4 and 5). Other details same as Fig. 2d.

In an exemplar participant, the burst % increased for Successful Stop compared to both Failed Stop and Correct Go trials prior to SSRT_Beh_ (Fig. 5a). To quantify this across participants, we compared the mean burst % among the 3 trial-types, and for the time window from the Stop signal to the SSRT_Beh_ of a participant (Stop_Win_) and the baseline period *before the Stop* signal (Base_Win_; Go to Stop signal in Stop trials and Go to mean SSD in Correct Go trials). We performed a 2-way repeated measures ANOVA with mean burst % as the dependent measure, with trial-type (Successful, Failed Stop, and Correct Go trials) and time-window (Stop_Win_ and Base_Win_) as factors. There was a significant main effect of trial-type (*F*(2,20) = 4.5, *p* = 0.025, *η_p_^2^* = 0.3) and a trial-type by time-window interaction (*F*(2,20) = 4.0, *p* = 0.034 *η_p_^2^* = 0.3), but no main effect of time-window (*F*(1,10) = 3.8, *p* = 0.088, *η_p_^2^* = 0.3). Post hoc *t*-tests showed that in the Stop_Win_ there was a significant increase in burst % for Successful Stop (14.6±1.7 %) compared to both its baseline (9.9±1.7 %; *t*(10) = 3.3, *p_Bon_* = 0.022, *BF*_10_ = 7.6), and Correct Go (9.6±1.3 %; *t*(10) = 3.7, *p_Bon_*= 0.015, *BF*_10_ = 11.8), but not to Failed Stop (10.3±1.6 %; *t*(10) = 2.1, *p_Bon_*= 0.198, *BF*_10_ = 1.2) (Fig. 5b). Thus, burst % increased for the Successful Stop trials.

To further clarify the temporal relationship between beta activity and our EMG measure of action-stopping, we quantified the mean burst time (BurstTime in the Stop_Win_) for each participant. Across participants, the mean BurstTime (115±6 ms) was significantly shorter than mean CancelTime (169±10 ms; *t*(10) = 8.2, *p* < 0.001, *BF*_10_ > 100) and there was also a strong positive relationship between them (*ρ* = 0.76, *p* = 0.006, *BF*_10_ = 10.6; Fig. 5d) [see Supplementary Fig. 5 for correlation between CancelTime and other burst parameters]. Further, we show that the observed correlation was not merely an artifact of varying Stop_Win_ across participants (permutation test, *p* < 0.05; see Methods). Thus, these results show that participants with an early frontal beta burst also had an early CancelTime.

### Study 5 (EEG replication)

We ran a new sample of 13 participants (see Table 1 for behavioral results). As above a right frontal IC was extracted for each participant (average topography Fig. 5c *inset*, see Supplementary Fig 3) and the burst % was compared for the 3 trial-types (Successful Stop, Failed Stop, and Correct Go) in the two time-windows (Stop_Win_ and Base_Win_). Again, a 2-way repeated measures ANOVA with burst % as the dependent measure revealed that there was a significant main effect of trial-type (*F*(2,24) = 6.9, *p* = 0.004, *η_p_^2^* = 0.4) and a trial-type by time-window interaction (*F*(1,12) = 5.8, *p* = 0.009, *η_p_^2^* = 0.3; Fig. 5c). Here there was also a significant effect of time-window on burst % (*F*(1,12) = 16.1, *p* = 0.002, *η_p_^2^* = 0.6). Post-hoc *t*-tests confirmed that the burst % was greater for Successful Stop (16.2±2.2 %) compared to its baseline (11.3±1.4 %; *t*(12) = 3.3, *p_Bon_* = 0.021, *BF*_10_ = 7.6), and Correct Go (12.0±1.4 %; *t*(12) = 3.0, *p_Bon_* = 0.030, *BF*_10_ = 5.3) but not compared to Failed Stop (15.4±1.4 %; *t*(12) = 1.0, *p_Bon_* = 0.957, *BF*_10_ = 0.34). Across participants, the mean BurstTime (129±7 ms) was again significantly shorter than CancelTime (166±8 ms; *t*(10) = 5.0, *p* < 0.001, *BF*_10_ > 100) and there was a significant positive relationship (*ρ* = 0.57, *p* = 0.045, *BF*_10_ = 1.9; Fig. 5d). Again, a permutation test suggested that this correlation was unlikely to result from mere variation in the length of StopWin across participants (*p* < 0.05). Combining data from studies 4 and 5 confirms the strong relationship between right frontal beta BurstTime and CancelTime (*ρ* = 0.66, *p* < 0.001, *BF*_10_ = 29.4).

## Discussion

This set of studies provides detailed information about the timing of subprocesses in human action-stopping. We started with the recently published observations that the standard behavioral measure of action-stopping (SSRT) is, an over-estimate of stopping speed^15,21,22^. To more precisely delve into this, we developed and validated a trial-by-trial method for estimating stopping speed from EMG. We focused on Successful Stop trials with small impulses (partial bursts) in EMG activity. The amplitude of such partial EMG activity was ∼50% of the amplitude of EMG activity for outright keypresses, and this decreased at ∼160 ms after the Stop signal (CancelTime). While, one interpretation of this partial EMG activity is that it merely reflects ‘weak’ Go activation that did not run to completion, several lines of evidence strongly suggest it is a muscle manifestation of the stopped response. First, CancelTime had a strong positive correlation with SSRT_Beh_. Second, the variability of CancelTime was positively correlated with the variability of SSRT estimated from the BEESTS modeling framework. Third, the partial EMG activity had a profile which was initially similar to the EMG profile seen when actual keypresses were made, and only diverged at ∼55 ms after EMG onset. This initial similarity would not be expected if it were a weak Go activation – since previous research has demonstrated that weak and strong muscle activations have distinct profiles that diverge soon after onset^39^. Fourth, our TMS experiment demonstrated that, CancelTime coincided well with the timing of a putative basal ganglia-mediated global motor suppression^25–30^. This implies that the smaller amplitude and earlier decline of the partial EMG activity on Successful Stop was due to an active suppression of motor output. Fifth, across participants, on Successful Stop trials, CancelTime correlated strongly with the time of right frontal beta bursts (BurstTime) from scalp EEG. This is consistent with response inhibition being implemented via right prefrontal cortex^12^, and with previous research showing an increase of beta at right frontal electrode sites before SSRT_Beh_^32,33^. Taken together, our results strongly suggest that CancelTime reflects the time of implementation of an active Stop process at the muscle-level. These results have striking theoretical and practical implications for response inhibition research and, more widely, our understanding of impulse control.

Notably, CancelTime was ∼60 ms earlier than SSRT_Beh_. To better understand this discrepancy, we calculated SSRT based on the EMG response rather than behavior. We saw that SSRT_EMG_ better matched CancelTime than did SSRT_Beh_. Thus, SSRT_Beh_ could be an over-estimation of the duration of the Stop process in the brain. This extra time in SSRT_Beh_ probably reflects a ‘ballistic stage’ in generation of the button press^40,41^. We suggest that the maximum CancelTime reflects the last point at which a Stop process can intervene to prevent responses. We note that CancelTime (a muscle measurement) is an overestimation of the brain’s stopping speed since it does not include the corticospinal conduction time, which we estimated at ∼20 ms. Indeed, our TMS results show that global motor suppression, which we take as the time at which motor areas of the brain are suppressed, is ∼140 ms (which is ∼15 ms less than CancelTime). One important consequence of our observation that the brain’s stopping speed is ∼140 ms is that neural events that mediate stopping need to occur before this time. Indeed, we found that right frontal beta activity increased ∼120 ms after the Stop signal on Successful Stop trials, and also that, across participants, there was a strong positive relationship between mean BurstTime and mean CancelTime.

Taken together, these studies motivate a detailed model of the temporal events of action-stopping (Fig. 6). First, we suppose the right frontal beta bursts relate to activity of right inferior frontal gyrus^12,31^, and this happens in ∼120 ms, which then leads via basal ganglia^29^ to global suppression of the primary motor cortex^25–28,30^ at ∼140 ms. After a corticospinal conduction delay of ∼20 ms, this suppression of motor output is then reflected at ∼160 ms as a decline in muscle activity (CancelTime). Finally, SSRT_Beh_ occurs at ∼220 ms, after, what we suppose is an electromechanical delay of ∼60 ms.

**Figure 6.**
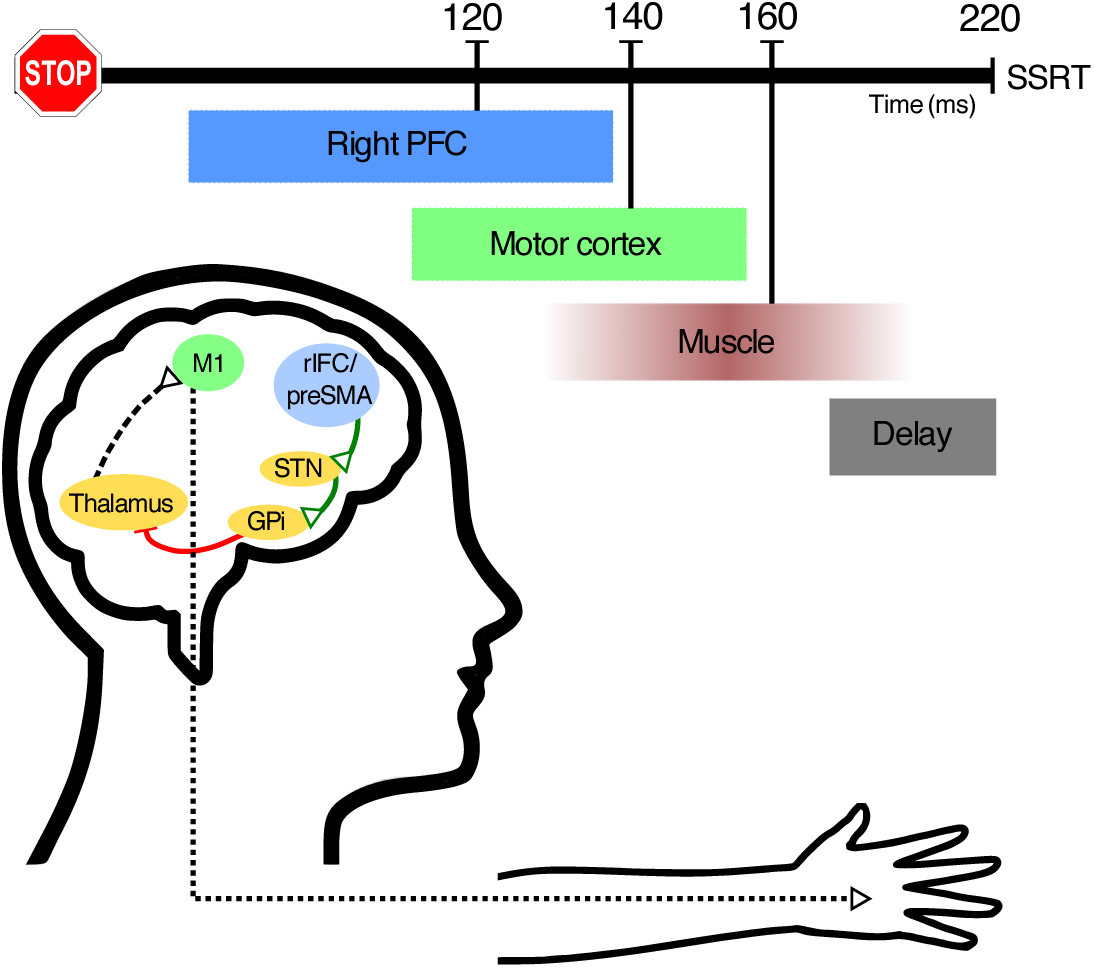
Hypothetical model of the temporal cascade of processes underlying human action-stopping. Following the Stop signal, the right PFC including the rIFC and the preSMA gets activated at ∼120 ms. These region/s activate the STN of the basal ganglia which in turn activates the globus pallidus interna which via its inhibition on the motor regions of the thalamus cuts down the ‘drive’ of the motor cortex. This results in a global motor suppression at ∼140 ms after the Stop signal. This suppression is reflected in the hand muscle at ∼160 ms which is measured as the CancelTime. There is a delay of ∼60 ms at the muscle level which gets added to the behavioral estimate of SSRT.

This model specifies the chronometrics of stopping in more detail than extant human models, and, more generally, raises questions about the timing reported in some other studies. For example, other research has shown that movement neurons in monkey Frontal Eye Field decrease activity <10 ms before SSRT^42^, dopaminergic neurons in rodent substantia nigra and striatum increase activity 12 ms prior to SSRT^43^, TMS at ∼25 ms before SSRT over human Intraparietal Sulcus prolongs SSRT^44^, and that P300 human EEG activity ∼300 ms after the Stop signal relates to stopping speed^45^. Our observations of shorter latencies for prefrontal bursts, TMS-MEP and muscle CancelTime, raise questions about what is reflected in these late neural activities.

Our results have several important implications. First, as just noted, they provide temporal constraints on neuroscience studies of stopping in the brain. They suggest that methods with high temporal resolution need to focus on the time after the Stop signal and before CancelTime (indeed CancelTime minus conduction time) rather than before SSRT_Beh_, and we provide a novel single-trial metric of stopping. Second, our results have clinical implications. Whereas meta-analysis shows that SSRT_Beh_ is longer for patients (e.g. ADHD, OCD, and substance use disorder) vs. controls^6–11^, not all such studies show differences^8,46–48^. We predict that our new single-trial method of CancelTime will be more sensitive than SSRT_Beh_. Furthermore, future studies can easily estimate within-subject variability in CancelTime, which will likely discriminate patients from controls. Third, our results provide insight into why SSRT_Beh_ might only have a modest relationship with more ‘real-world’ measures of impulsivity^15–20^. As we show, the SSRT_Beh_ includes not only CancelTime but an extra, and variable, 60 ms ballistic stage. We expect that future studies may show stronger correlations between CancelTime and self-report than that seen between SSRT_Beh_ and self-report (also see15); likewise we predict that right frontal beta burst time might also correlate more tightly with self-report measures. More generally, the detailed timing information of frontal beta at ∼120 ms, global motor suppression at ∼140 ms, and CancelTime at ∼160 ms points to subprocesses of action-stopping that provide potential biomarkers that could better explain individual differences in impulse control.

In conclusion, we provide a detailed timing model of action-stopping that partitions it into subprocesses that are isolable to different nodes and are surely more precise than the behavioral speed of stopping. At the core of this timing model is a novel method of measuring the speed of stopping from the muscles. This provides a single-trial estimate of stopping speed that could be easily measured with minimal equipment in any lab that studies human participants.

## Methods

### Participants

All were adult, healthy, human volunteers who provided written informed consent and were compensated at $20/hour. The studies were approved by the UCSD Institutional Review Board.

*Study 1*. Ten participants (4 females; age 22±1 years; all right-handed).

*Study 2*. Thirty-six participants (19 females; age 19±0.4 years; all right-handed). Two were excluded for bad behavior (violating the assumptions of the independent race model - Failed Stop RT < Correct Go RT, and P(Stop) increasing monotonically as a function of SSD), and two were excluded for noisy EMG data.

*Study 3 (TMS)*: Eighteen participants (11 females; age 19 ± 0.4 years; 15 right-handed, 2 left-handed) with no contraindications to TMS^49^. One was excluded for bad behavior.

*Study 4 (EEG)*. Eleven participants (6 females, age 19 ± 0.4 years, all right-handed) participated.

*Study 5 (EEG)*: Fifteen participants (9 females, age 21±0.4 years, all right-handed) participated. Two were excluded from analysis, one for misaligned EEG markers due to a technical issue, while the other lacked a right frontal brain IC, based on our standard method^32,33^.

### Stop-signal task

This was run with MATLAB 2014b (Mathworks, USA) and Psychtoolbox^50^. Each trial began with a white square appearing at the center of the screen for 500±50 ms. Then a right or left white arrow appeared at the center. When the left arrow appeared, participants had to press a key on a vertically oriented keypad using their index finger, while for a right arrow they had to press down on a key on a horizontally oriented keypad with their pinky finger (Fig. 1b *inset*), as fast and as accurately as possible (Go trials). The stimuli remained on the screen for 1s. If participants did not respond within this time, the trial aborted, and ‘Too Slow’ was presented. On 25% of the trials, the arrow turned red after a stop signal delay (SSD), and participants tried to stop the response (Stop trials). The SSD was adjusted using two independent staircases (for right and left directions), where the SSD increased and decreased by 50 ms following a Successful Stop and Failed Stop respectively. Each trial was followed by an inter-trial interval (ITI) and the entire duration of each trial including the ITI is 2.5 s (Fig. 1a).

*Study 1 and 2.* Participants performed the task with their right hand. They performed 40 practice trials before the actual experiment, where their baseline SSD was determined and was subsequently used as the starting SSD in the main experiment. In study 1 and 2, the experiment had 600 trials divided in 15 blocks, such that each block had 40 trials (450 Go trial and 150 Stop trials). At the end of each block the participants were presented a figure showing their mean reaction times (RT) in each block. Participants were verbally encouraged to maintain their mean reaction time constant across the different blocks and between 0.4 – 0.6 s.

*Study 3.* Participants performed the task with their left hand. Following 48 practice trials without TMS, participants performed 12 blocks of the experiment with TMS, with each block consisting of 96 trials each (72 Go trials and 24 Stop trials).

*Study 4.* Participants performed the task with their right hand. Following 160 practice trials, participants performed 4 blocks of 80 trials (240 Go trials and 80 Stop trials).

*Study 5.* Participants performed the task with their right hand. Following 80 practice trials, participants 24 blocks of 80 trials each (1440 Go trials and 480 Stop trials).

### EMG recording

EMG data were acquired using a Grass QP511 AC amplifier (Glass Technologies, West Warwick, RI) with a frequency cut-off between 30 and 1000 Hz. A CED Micro 1401 mk II acquisition system sampled the data at 2 kHz. The EMG data were acquired by CED Signal v4 software (Cambridge Electronic Design Limited, Cambridge, UK) for 2 s following the fixation cue. The data acquisition was triggered from MATLAB using a USB-1208FS DAQ card (Measuring Computing, Norton, MA). In all 5 experiments, surface EMG was recorded from both the first dorsal interossei (FDI) and the abductor digiti minimi (ADM) muscles of the hand (Fig. 1b *inset*). In the TMS experiment, surface EMG was also recorded from the task-irrelevant right extensor carpi radialis (ECR) muscle (Fig. 5a).

### TMS

MEPs were evoked using a TMS device (PowerMag Lab 100, MAG&More GMBH, Munich, Germany) delivering full sine wave pulses, and connected to a figure-of-eight coil (70 mm diameter, Double coil PMD70-pCool; MAG&More GMBH, Munich, Germany). During the task, the coil was positioned on the scalp over the left primary motor cortex representation of the ECR muscle and oriented so that the coil handle was approximately perpendicular to the central sulcus, *i.e.* at ∼45° to the mid-sagittal line, and the initial phase of current induced in the brain was posterior-to-anterior across the central sulcus. Prior to the experiment, the motor hot spot was determined as the position on the scalp where slightly supra-threshold stimuli produced the largest and most consistent MEPs in ECR. The position was marked on a cap worn by the participants. Resting motor threshold (RMT) was defined as the lowest intensity to evoke an MEP of at least 0.05 mV in 5 of 10 consecutive trials while participants were at rest. We then established the test stimulus intensity to be used during task, which was set to produce a mean MEP amplitude of approximately 0.2 - 0.5 mV whilst the participant was at rest.

MEPs were also evoked in the left FDI muscle prior to beginning the main experiment for the purpose of recording the corticospinal conduction time. The motor hot spot for the FDI was defined in a manner similar to that for the ECR. The active motor threshold (AMT) was defined as the lowest intensity to evoke a discernible MEP in 5 of 10 consecutive trials, while participants maintained slight voluntary contraction (∼10% of maximum voluntary EMG amplitude during isometric finger abduction). Then, 10 stimuli were delivered at 150% AMT during slight voluntary contraction (again 10% of maximum), with the coil oriented to induce lateral-medial current in the brain in order to obtain estimates of corticospinal conduction time.

During the task, TMS stimuli were delivered on every Stop trial and on 50% of Go trials. On every Stop trial, a single TMS stimulus at the test stimulus intensity was delivered at one of six time points: inter-trial interval (100 ms prior to fixation; ITI), 100 ms, 120 ms, 140 ms, 160 ms and 180 ms after the Stop signal (Fig. 4a). On the Go trials, TMS stimuli were yoked to the time of the Stop signal on the previous Stop trial. Thus, there were 48 trials per TMS time point on Stop trials and 96 trials per time point on Go trials.

### EEG

64 channel EEG (Easycap, Brainvision LLC) was recorded in the standard 10/20 configuration at 1 kHz.

### Data analysis

All analyses were performed using MATLAB (R2016b, R2018b, R2019a).

#### Stop Signal Reaction Time

SSRT from the behavioral responses (SSRT_Beh_) was determined using the integration method5. When calculating SSRT using the EMG responses, SSRT_EMG_, as the P(Respond|Stop) was often much more than 0.5, we calculated the SSRT individually for the 3 most frequent SSDs and then averaged it^51^.

#### EMG data analysis

EMG data were filtered using 4^th^ order Butterworth filter (roll-off 24 dB/octave) to remove 60 Hz noise and its harmonics at 120, and 180 Hz. EMG data were full-wave rectified and the root-mean square (RMS) of the signal was computed using a centered window of 50 ms. Any EMG activity which was greater than 8 SD of the mean EMG activity in the baseline period (Fixation to Go cue) was marked, on a trial-by-trial basis. Starting from the peak of that EMG activity, the onset was marked at the point where the activity dropped below 20% of the peak for 5 consecutive ms. This method of adjusting the threshold based on the peak EMG activity, allowed better onset detection than a fixed threshold, especially when the amplitude of the EMG activity was small. The time when EMG started to decline was determined as the time when, following the peak EMG activity, the activity decreased for 5 consecutive ms. Visual inspection of individual trials showed that this method provided a reliable detection of both EMG onsets (see Supplementary Fig. 1a, 1b for EMG onset vs. RT correlation) and decline. Any detected EMG timing which was beyond 1.5 times the inter-quartile range (IQR) of the first and third quartile (Q3) of that particular timing distribution was deemed an outlier. This removed <4% trials. CancelTime was marked as the time of the decline following the Stop signal. For outlier rejection, CancelTimes had a lower cutoff of 50 ms and higher cutoff of Q3+1.5×IQR. This removed <3% trials.

As the peak EMG amplitude for the FDI and ADM muscle were quite distinct, before averaging the two EMG activities, we normalized the muscle activity by the peak activity in that particular muscle (Voltage_Norm_ in Fig. 2a, 2b, 3c, Supplementary Fig. 1c).

#### Global MEP suppression

MEP amplitudes were measured on a trial-by-trial basis. Data were included for analysis if the following criteria were met: *(i)* the amplitude of the ECR EMG signal in a 90 ms period prior to the TMS stimulus was < 0.05 mV; *(ii)* the amplitude of the MEP fell within the mean±1.5×IQR of values for the same time point and trial type (Correct Go, Failed Stop, Successful Stop). Thereafter, MEP amplitudes measured at the ITI were collapsed across trial type (Correct Go, Failed Stop and Successful Stop), averaged and used as a baseline against which to compare other TMS time points. For each of the other TMS time points (100, 120, 140, 160, 180 ms following the Stop signal), data were averaged within each trial type (Correct Go, Failed stop, Successful Stop) and expressed as a percentage of the mean ITI MEP amplitude.

#### Corticospinal conduction time

Corticospinal conduction time was determined by delivering TMS over the hand representation of left FDI and measuring MEP from the muscle (Fig. 4c). The earliest MEP onset latency across 10 trials was identified by visual inspection of the EMG traces^52–54^.

#### Trial-by-trial analysis of the time of the CancelTime and time of global motor suppression

To compare the temporal association between the EMG decline and MEP suppression, we performed a trial-by-trial analysis of stop-signal task data only on trials where an EMG burst was detected. We first normalized the time of TMS on a given trial by subtracting the time of EMG decline from the time of the TMS pulse. Hence, negative values mean that TMS was delivered before the EMG decline and positive values mean that TMS was delivered after. We then plotted MEP amplitudes for each of the three response types (Correct Go, Failed Stop, and Successful Stop) against the normalized times binned into 30 ms windows. This analysis meant that for a given individual there were relatively few trials per time bin, and some bins would occasionally contain no data. Therefore, we combined data across all individuals. Prior to this, MEP amplitudes for each individual were normalized to the mean MEP amplitude at the inter-trial interval, to account for inter-individual variability in absolute MEP amplitudes at baseline. We restricted our analysis to time bins that contained at least 50 trials, which resulted in time range −90 ms to 60 ms.

#### EEG Preprocessing

We used EEGLAB^55^ and custom-made scripts to analyze the data. The data were downsampled to 512 Hz and band-pass filtered between 2-100 Hz. A 60 Hz and 180 Hz FIR notch filter were applied to remove line noise and its harmonics. EEG data were then re-referenced to the average. The continuous data were visually inspected to remove bad channels and noisy stretches.

#### ICA analysis

The noise-rejected data were then subjected to logistic Infomax ICA to isolate independent components (ICs) for each participant separately^38^. We then computed the best-fitting single equivalent dipole matched to the scalp projection for each IC using the DIPFIT toolbox in EEGLAB^55,56^. ICs representing non-brain activity related to eye movements, muscle, and other sources were first identified using the frequency spectrum (increased power at high frequencies), scalp maps (activity outside the brain) and the residual variance of the dipole (greater than 15%) and then, subtracted from the data. A putative right frontal IC was then identified from the scalp maps (if not present then we used frontal topography) and the channel data were projected onto the corresponding right frontal IC. The data on Successful Stop trials were then epoched from −1.5 s to 1.5 s aligned to the Stop signal. We estimated the time-frequency maps from 4 to 30 Hz, and −100 to 400 ms using Morlet wavelets with 3 cycles at low frequencies linearly increasing by 0.5 at higher frequencies. The IC was selected only if there was a beta power (13 to 30 Hz) increase in the window between the Stop signal and SSRT_Beh_ compared to a time-window *prior to the Go* cue (−1000 to −500 ms aligned to Stop signal). In each participant, the beta frequency which had the maximum power in this time window was used in the beta bursts computation (Supplementary Fig. 3).

#### Beta Bursts

To estimate the beta bursts, the epoched data were first filtered at the peak beta frequency using a frequency domain Gaussian window with full-width half-maximum of 5 Hz. The complex analytic envelope was then obtained by Hilbert transform, and its absolute value provided the power estimate. In each participant, to define the burst threshold, the beta amplitude within a period of 500 to 1000 ms (*i.e.* after the Stop signal in the Stop trials, and after the mean SSD in the Correct Go trials) was pooled across all trials [compared to the ICA analysis here we picked a different time-window to estimate the burst threshold to keep the analysis unbiased. However, picking the same time-window also yielded similar results]. The threshold was set as the median + 1.5 SD of the beta amplitude distribution (Supplementary Fig. 4). Once the burst was detected, the burst width threshold was set as the median + 1 SD. We binary-coded each time point where the beta amplitude crossed the burst width threshold to compute the burst % across trials. For each detected burst, the time of the peak beta amplitude was marked as the BurstTime.

### Statistical analysis

For pairwise comparisons, the data were first checked for normality using Lilliefors test, and if normally distributed a two-tailed *t*-test (*t*-statistic) was performed, else a Wilcoxon signed rank test (*Z*-statistic) was performed. We interpret the effect sizes as small (Cohen’s *d*: 0.2-0.5; Bayes Factor in favor of the alternate hypothesis, *BF_10_*: 1-3), medium (*d*: 0.5-0.8; *BF_10_*: 3-10), large (*d* > 0.8; *BF_10_* > 10). For comparisons across multiple levels, repeated-measures ANOVA was used, followed by Bonferroni corrected *t*-tests for pairwise comparisons (Bonferroni corrected *p*-value: *p_Bon_*). The Greenhouse-Geisser correction was applied where the assumption of sphericity in ANOVA was violated (corrected *F*-statistic: *FGG*). Effect sizes for ANOVAs were interpreted as small (partial eta-squared, *η_p_^2^*: 0.01-0.06), medium (*η_p_^2^*: 0.06-0.14), and large (*η_p_^2^* > 0.14). For correlational analyses, Pearson’s correlation coefficient (*r*) was usually used, but Spearman’s correlation coefficient (*ρ*) was used when the data bounded in a closed interval. All data are presented as mean±s.e.m.

In testing the relationship between BurstTime and CancelTime, we performed a permutation test. We sampled BurstTimes randomly from a uniform distribution between 0 and SSRT_Beh_ for a given participant for 3000 iterations. For each iteration, we then computed the correlation (*r*) between the mean BurstTime and the mean CancelTime across participants. This generated a distribution of *r* ranging between −1 and 1. The *p*-value for our analysis was determined as the P(*r*≥*r_Obs_*|H_0_) in the permuted data.

## Bayesian modelling of behavioral data

We used the BEESTs model developed by Dora Matzke and colleagues (run in R Studio 1.1.463) which assumes a race between two stochastically independent process, a Go and a Stop processes. This model estimates the distribution of the SSRT by using the participant’s Go RT distribution, and by considering the Failed Stop RTs as a censored Go RT distribution. The censoring points are sampled randomly from the SSRT distribution on each Stop trial. The RT distributions underlying the Go and Stop process is assumed to have a Gaussian and an exponential component and is described 3 parameters (*μ_Go_*, *σ_Go_*, *τ_Go_* and *μ_Stop_*, *σ_Stop_ τ_Stop_*). For such ex-Gaussian distributions, the mean and variance of the RT distribution are determined as *μ* + *τ* and *μ*^2^ + *τ*^2^, respectively. The model also estimates the probability of trigger failures for each participant. The model uses Bayesian Parametric Method (BPE) to estimate the parameters of the distributions. We used a hierarchical BPE, where individual subject parameters are modeled with the group-level distributions. This approach is thought to be more accurate than fitting individual participants and is effective when there is less data per participant57. We pooled the subjects across both study 1 and 2 to estimate the individual parameters. The priors were bounded uniform distributions (*μ_Go_*, *μ_Stop_*: *U*(0,2); *σ_Go_*, *σ_Stop_*: *U*(0,0.5) *τ_Go_*, *τ_Stop_*: *U*(0,0.5); pTF: U(0,1)). The posterior distributions were estimated using the Metropolis-within-Gibbs sampling and we ran multiple chains. We ran the model for 5000 samples with a thinning of 5. The Gelman-Rubin (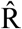) statistic was used to estimate the convergence of the chain. Chains were considered converged if 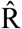 < 1.1.

## Acknowledgements

We thank Dora Matkze for sharing the scripts for BEESTS modelling, Sven Bestmann for insightful comments on data, Simon Little for sharing the beta-burst analysis script, Kelsey Sundby for sharing some EEG and EMG data, and Xinze Yu and Hunter Robbins for help in data recording. We gratefully acknowledge our support from NIH: NS106822 and DA026452.

## Author contributions

S.J., R.H., V.M., and A.R.A conceived the experiments; S.J., R.H., and V.M. recorded and analyzed the data; S.J., R.H., V.M., and A.R.A wrote the paper.

## Data and scripts

A core element of this paper is a novel method of calculating single-trial stopping speed from EMG. Accordingly, we provide the EMG and behavioral data from 10 participants in study 1, along with analysis scripts, and a brief description of how to execute the scripts (https://osf.io/b2ng5/). Upon acceptance of the paper all EMG, TMS-MEP and EEG data will be provided at the above link.

