## Supplementary Figures for "Temporal cascade of frontal, motor and muscle processes underlying human action-stopping"

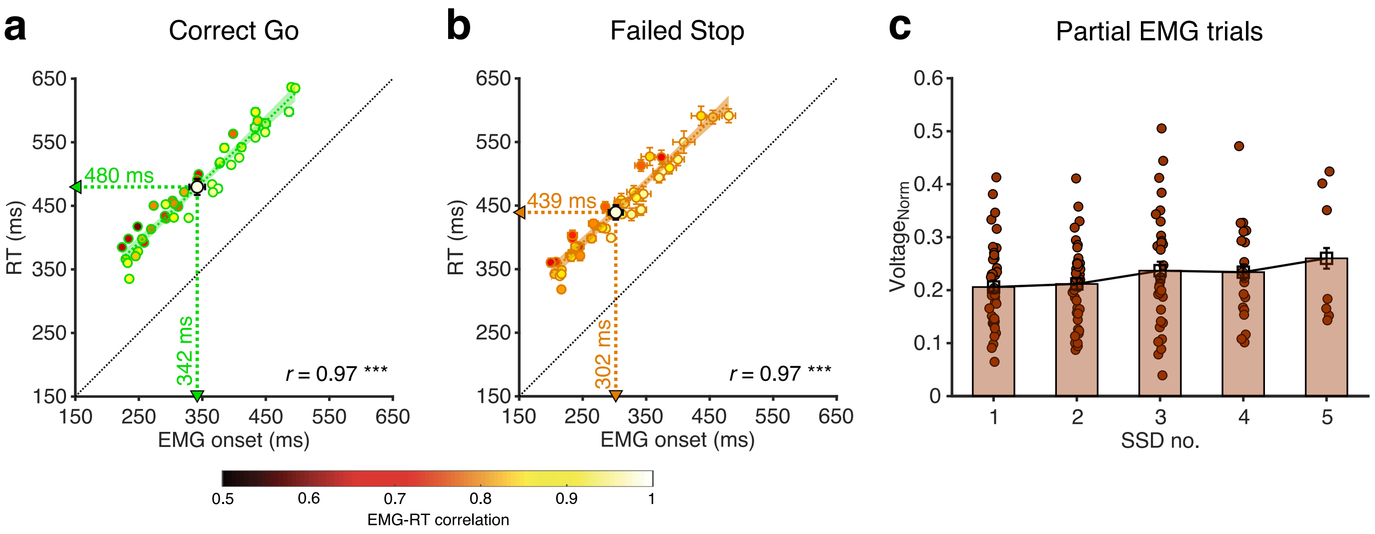


**Supplementary Figure 1 I EMG responses in study 1 and 2.** (**a**) EMG onset-RT correlation in the Correct Go trials. Each small dot represents the mean in a participant, color coded by the EMG-RT correlation in that participant, while the cross-hairs represent the s.e.m. in that participant. The large dot and cross-hairs represent the mean and s.e.m. across all the participants, again color coded by the mean EMG-RT correlation ($r = 0.97$, $p < 0.001$, $BF_{10} > 100$; EMG onset = 342±13 ms, RT = 480±12 ms) across all participants. The linear regression fit and its 95% confidence interval is shown as a green line and shaded region respectively. The unity line is represented as a black dotted line. (**b**) Same as (a) but for the Failed Stop trials ($r = 0.97$, $p < 0.001$, $BF_{10} > 100$; EMG onset = 302±11 ms, RT = 439±11 ms). (**c**) The normalized EMG voltage in the partial EMG trials as a function of the first 5 SSDs (SSD no.) in each participant ($SSD_1$ = 0.206±0.012, $SSD_2$ = 0.212±0.012, $SSD_3$ = 0.237±0.017, $SSD_4$ = 0.234±0.015, $SSD_5$ = 0.260±0.020; rmANOVA: $F(4,24) = 3.7$, $p = 0.018$, $\eta_p^2 = 0.4$). Each dot represents a participant, while the bar and cross-hairs represent the mean and s.e.m. across all the participants.

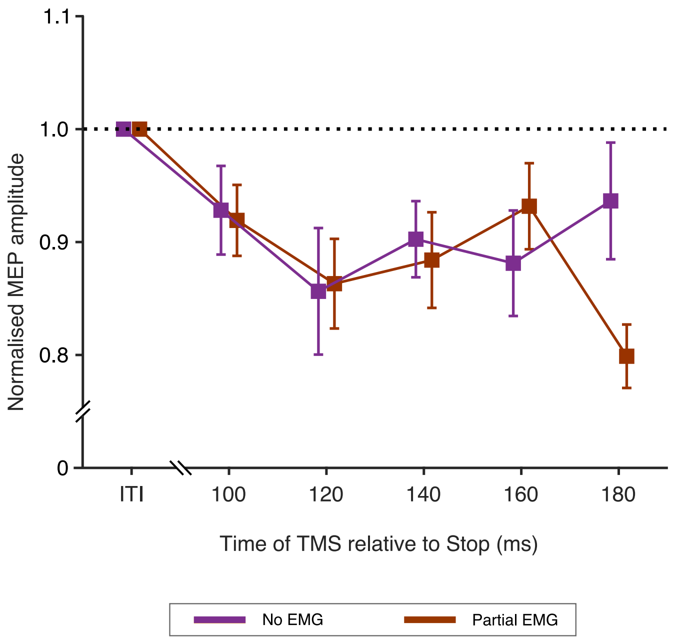


**Supplementary Figure 2 l MEPs in study 3.** Normalised MEP amplitudes in Successful Stop trials divided according to the presence, or not, of a partial EMG burst. There was no difference in amplitude of MEPs for trials with (Partial EMG) and without (No EMG) EMG bursts. rmANOVA showed no main effect of trial type ($F(1,16) = 1.2$, $p = 0.288$, $\eta_p^2 = 0.1$) or time ($F(4, 64) = 1.1$, $p = 0.356$, $\eta_p^2 = 0.1$), and interaction of trial-type by time ($F(4, 64) = 2.3$, $p = 0.066$, $\eta_p^2 = 0.1$). Therefore the timing and extent of the global motor system suppression on Successful Stop trials was not influenced by whether or not a partial EMG burst was detected.

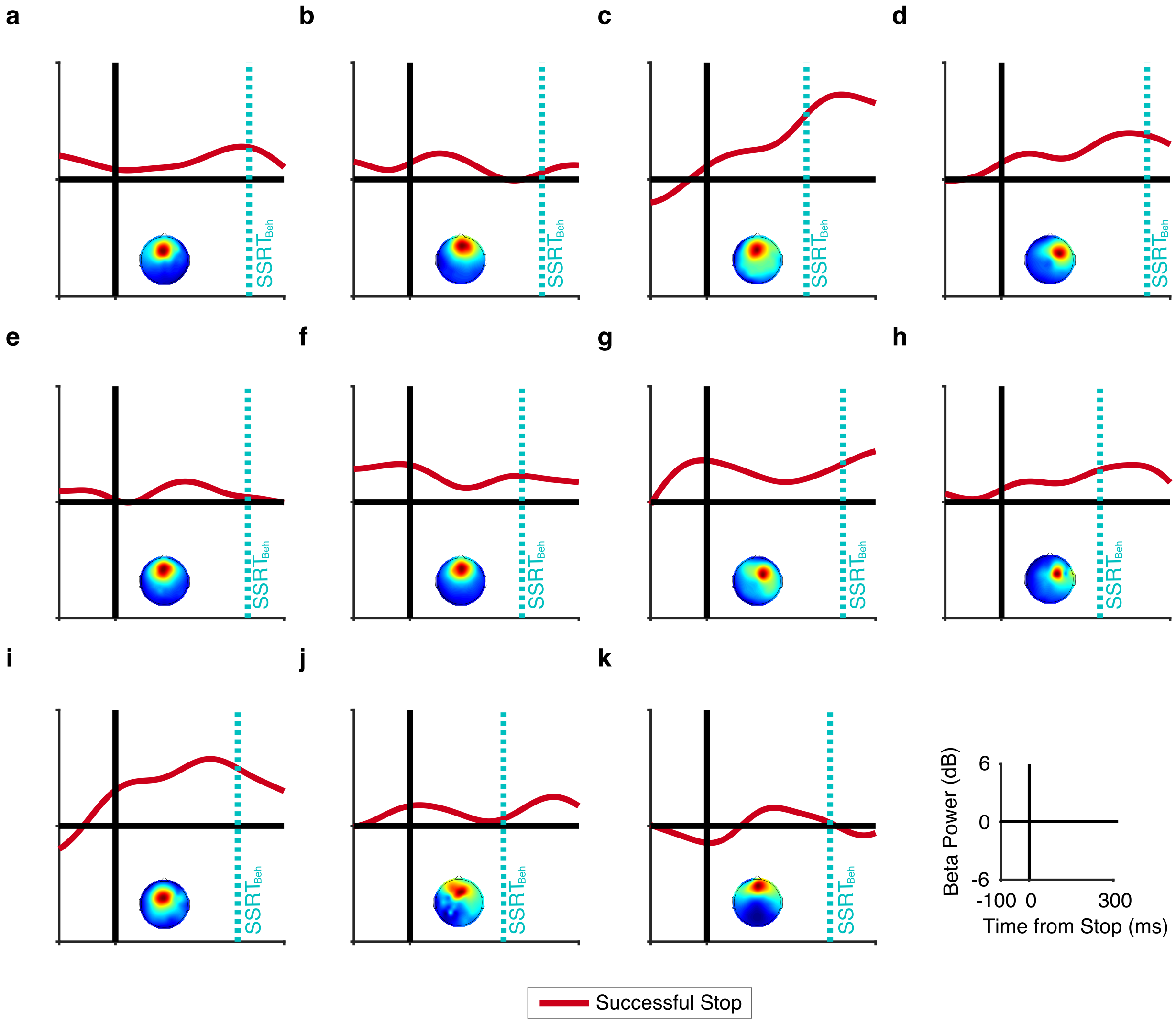


**Supplementary Figure 3 I Beta power in Successful Stop trials for study 4.** (**a-k**) shows the time course of the peak beta power (solid red line) in the selected IC for each participant. The dotted cyan line represents each participant's $SSRT_{Beh}$ . The *inset* in each panel shows the scalp topography for the selected IC.

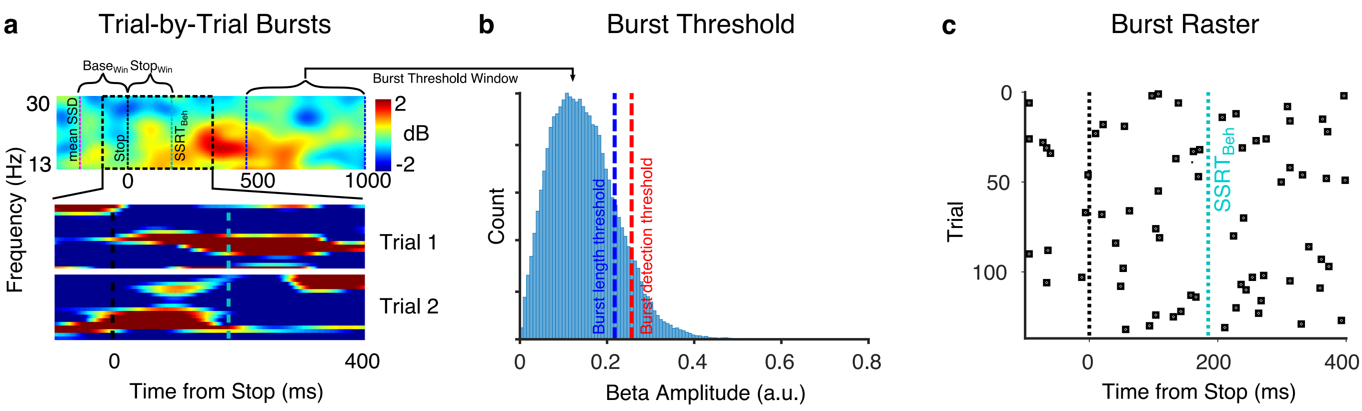


**Supplementary Figure 4 I Illustration of beta burst computation.** (**a**) The average beta power (13-30 Hz) between the Stop signal and $SSRT_{Beh}$ (top panel) for a participant. The window between mean SSD (dotted pink line) and Stop (dotted black line) represents the baseline time window ($Base_{Win}$). The time from Stop signal to $SSRT_{Beh}$ (dotted cyan line) represents the $Stop_{Win}$. The burst threshold window is the time period from 500 - 1000 ms after the Stop Signal. The bottom two panels show single trial examples of brief bursts of activity with the dotted black and cyan lines represent the Stop signal and $SSRT_{Beh}$ respectively. (**b**) The histogram of beta amplitude for all trials in a participant (500 to 1000 ms after the Stop signal in the Stop trials; 500 to 1000 ms after mean SSD in the Go trials). The dotted red line represents the threshold for defining a burst which is estimated as median + 1.5 SD of the beta amplitude. The BurstTime and burst height are the time and amplitude of the peak within a burst respectively. Once a burst is identified using this threshold, the burst % and burst length are estimated using a lower threshold which is median + SD (dotted blue line) (**c**) The burst raster for all trials in a participant across time. Each black dot represents a burst in that trial. To compute BurstTime we consider the burst times between the Stop signal and $SSRT_{Beh}$ (dotted cyan line).

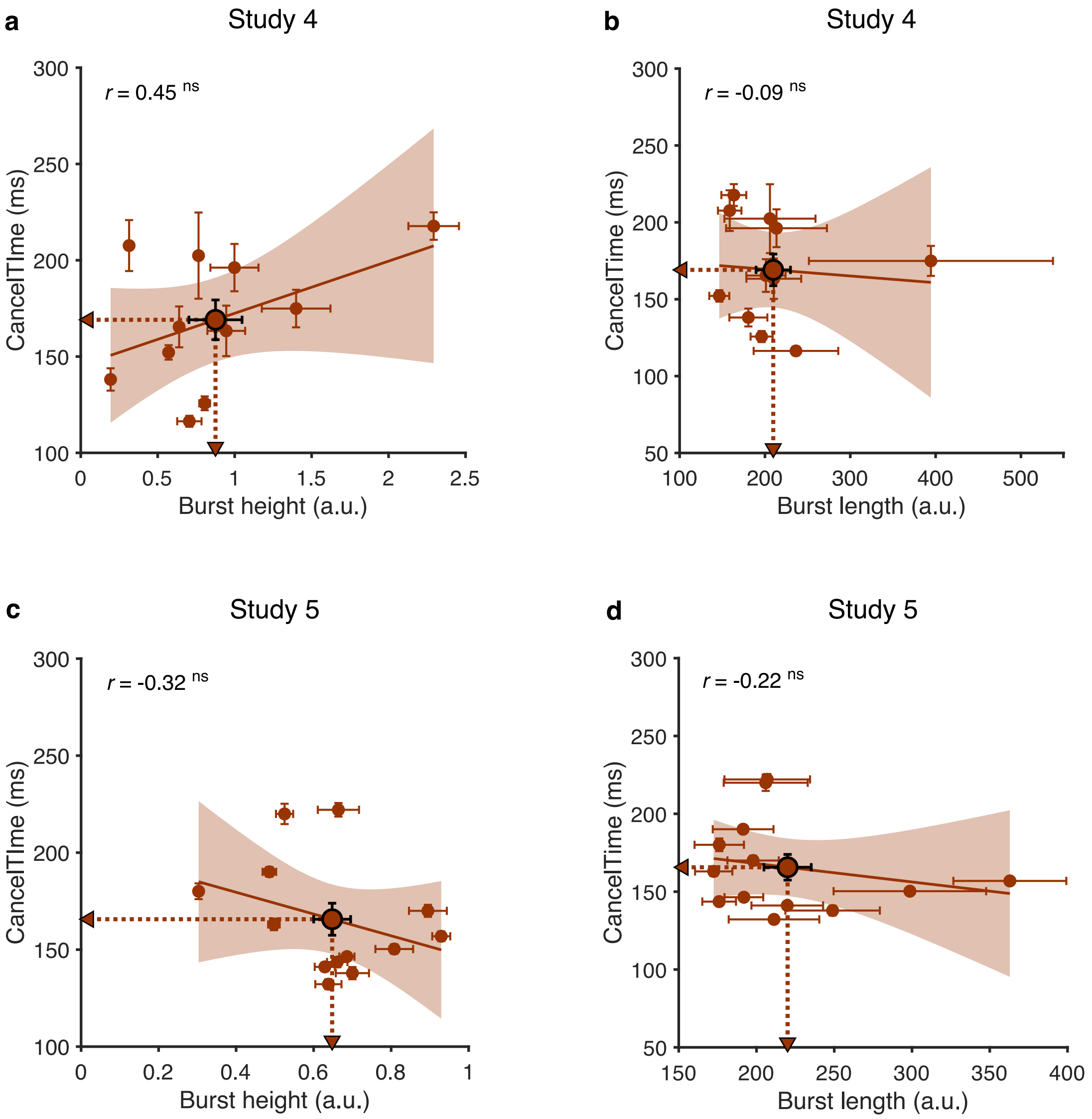


**Supplementary Figure 5 | Relationship of CancelTime with other burst parameters.** (**a**) Correlation between mean burst height (0.9±0.2) and mean CancelTime across participants in study 4 ($r$ = 0.45, $p$ = 0.161, $BF_{10}$ = 0.7). The brown dots represents individual participants and the cross-hairs represent the s.e.m. of the respective variables. The brown line and the shaded area represents the linear regression fit and its 95% confidence interval respectively. (**b**) Across participants correlation between mean burst length (210±20 ms) and mean CancelTime in study 4 ($r$ = -0.09, $p$ = 0.801, $BF_{10}$ = 0.3). Other details are same as in (a). (**c**) Same as (a) but for study 5. Mean burst height is 0.65±0.05 ($r$ = -0.32, $p$ = 0.281, $BF_{10}$ = 0.5). (**d**) Same as (b) but for study 5. Mean burst length is 220±15 ms ($r$ = -0.22, $p$ = 0.475, $BF_{10}$ = 0.4).
